# Systematic benchmarking of multi-modal approaches for tumor-naïve ctDNA detection and quantification

**DOI:** 10.64898/2026.06.19.733293

**Authors:** Ting Qi, Denis Odinokov, Lakshmi Narayanan Lakshmanan, Nathalia Graf Grachet, Melanie Lou, Seng Saelee, Gladys Garcia-Montoya, Wong Pui Mun, Rafeed Chowdhury Rahman, Hosseinali Asgharian, Andrea Teh Xin Yi, Nicole Hnin Ye Pyone, Lan Ying Wang, Guat Tin Tan, Hanaé Carrié, Avril Lim, Lau Yi Ting, Anna Gan Hwee Hsia, Polly Poon Suk Yean, Steven Ngo, Jordan Snyder, Harpreet Kaur, Aaron Tan, Yoon Sim Yap, Daniel SW Tan, Iain Bee Huat Tan, Jo-Anne Penkler, Sowmi Utiramerur, Dinesh Kumar, Anders Jacobsen Skanderup

## Abstract

Longitudinal monitoring of circulating tumor DNA (ctDNA) has emerged as a promising framework for characterizing treatment response dynamics in cancer. Scalable tumor-naïve approaches for quantifying ctDNA often involve whole-genome sequencing (WGS) or DNA methylation profiling, but their comparative performance and capacity for complementary integration remain poorly understood. Here we systematically benchmarked tumor-naïve WGS- and methylation-based ctDNA quantification methods using plasma from 150 patients with colorectal, lung and breast cancer. Using paired high-depth WGS and EM-seq data, we generated 40,000 in silico samples and evaluated detection accuracy, limits of detection (LoD) and quantification (LoQ) across cancer types and sequencing depths (0.1x–30x). We further assessed single- and multimodal method combinations, identifying conditions under which integrated approaches enhance analytical performance for detection and quantification relative to single modalities. This benchmark delineates key performance trade-offs and provides a practical framework to support method development and guide future research applications in ctDNA-based biomarker studies.

## Introduction

Circulating cell-free DNA (cfDNA) has emerged as a promising biomarker for non-invasive disease detection and monitoring. In healthy individuals, plasma cfDNA derives largely from turnover of hematopoietic cells, whereas in cancer patients it also contains circulating tumor-derived DNA fragments (ctDNA) (1,2). The ctDNA abundance provides a non-invasive proxy for the tumor burden and underpins a broad range of clinical applications, including cancer detection, enhanced prognostic staging, treatment monitoring, and minimal residual disease assessment (3–7). Indeed, accumulating evidence indicates that ctDNA kinetics serves as a robust early predictor of treatment response across diverse therapeutic modalities, including cytotoxic, targeted, and immunotherapies (8–12).

Molecular profiling using low-pass WGS (lpWGS) has emerged as a scalable tumor-naïve approach for ctDNA detection and quantification (13–16). This approach requires low blood volumes, involves simple library preparation, eliminates the need for tumor tissue, and reduces sequencing cost. Methods based on lpWGS exploit distinct genome-wide molecular features in ctDNA, such as DNA methylation signatures (17), copy number alterations (CNAs) (13), or fragmentation patterns (16,18). Numerous computational methods have been developed to estimate ctDNA levels from these genome-wide signals. Approaches based on CNAs or fragmentomics are often tumor-agnostic, whereas methylation-based methods typically rely on markers predefined for individual cancer types. Yet, limits of detection (LoD) and limits of quantification (LoQ) are not consistently reported, and direct comparisons across methods, cancer types, and sequencing depths remain scarce.

Existing method benchmarks have primarily focused on DNA methylation-based cell-type and tissue deconvolution rather than on ctDNA detection and quantification (19,20). While one study evaluated both cell-type deconvolution and ctDNA detection in cancer patients (20), it did not assess limits of detection and compare performance with WGS-based methods in cancer patients. A comprehensive study previously compared WGS and DNA methylation sequencing approaches for cancer detection in a large cohort of cancer and healthy patients (21). However, this study focused on proprietary methods and did not include publicly available tools; moreover, neither the datasets nor the methods are accessible, limiting its value as a community benchmark. To date, no study has systematically evaluated publicly available methods for ctDNA detection and quantification across data modalities, sequencing depths, and cancer types.

Here, we present a systematic benchmark of WGS-based methods for estimating ctDNA levels (tumor fraction, TF) in plasma samples. We generated matched WGS (∼50x coverage) and whole-genome DNA methylation sequencing via enzymatic conversion (EM-seq, ∼50x) data from the same plasma specimens obtained from 150 patients with colorectal, lung, or breast cancer, as well as 16 healthy individuals. We surveyed the published literature and identified 29 potential methods, of which 8 provided publicly available code and marker sets for evaluation. We constructed over 40,000 in silico samples and evaluated each method’s detection accuracy, quantification concordance, limit of detection (LoD) and quantification (LoQ) across cancer types and sequencing depths of 30x, 5x, 1x, and 0.1x. We further assessed multi-modal combinations of WGS-and methylation-based methods to identify conditions under which they improve sensitivity and accuracy.

## Results

### Overview of benchmark approach

We performed an exhaustive literature survey to identify computational methods capable of quantifying ctDNA levels (tumor fraction, TF) from whole-genome cfDNA sequencing (WGS or EM-seq). We initially identified 29 candidate methods based on distinct cfDNA-derived signals, including copy-number alterations (CNAs), fragmentomic features, and DNA methylation signatures (Methods; Suppl. Table S1). We then restricted the set to methods with publicly available, complete software packages that could be executed on new data. For DNA methylation-based approaches, we additionally required the availability of the methylation marker sets necessary for deconvolution. Finally, we performed software sanity checks to confirm that each method produced valid and interpretable outputs. These steps yielded eight methods for subsequent benchmarking: two based on WGS data (Fragle (16) and ichorCNA (13)) and six based on EM-seq data (cfNOMe (22), cfSort (23), MethAtlas (1), MIMESIS (24), SRFD-Bayes (25), and UXM (26)).

To establish an independent multi-cancer benchmark cohort, we collected plasma from 150 patients with colorectal, lung or breast cancer, as well as from 16 healthy individuals (Fig. 1b, Suppl. Table S2). We generated and processed deep whole-genome sequencing (median depth 78x) and whole-genome Enzymatic Methyl-seq (EM-seq; median depth 84x) data for all samples (Methods; Suppl. Fig. S1). Consensus TFs were estimated for each sample by taking the median across methods, and all cancer-type cohorts included samples with both low and high TFs, as inferred by the methods (Fig. 1). From these datasets, we generated down-sampled technical replicates (30x, 5x, 1x and 0.1x) and in silico dilution series by mixing cancer and healthy samples in controlled proportions (down to 0.5% TF). In total, this approach generated 40,160 down-sampled and in silico-diluted samples, which were used to evaluate each method’s detection accuracy, quantitative concordance, and limits of detection (LoD) and quantification (LoQ).

**Figure 1:**
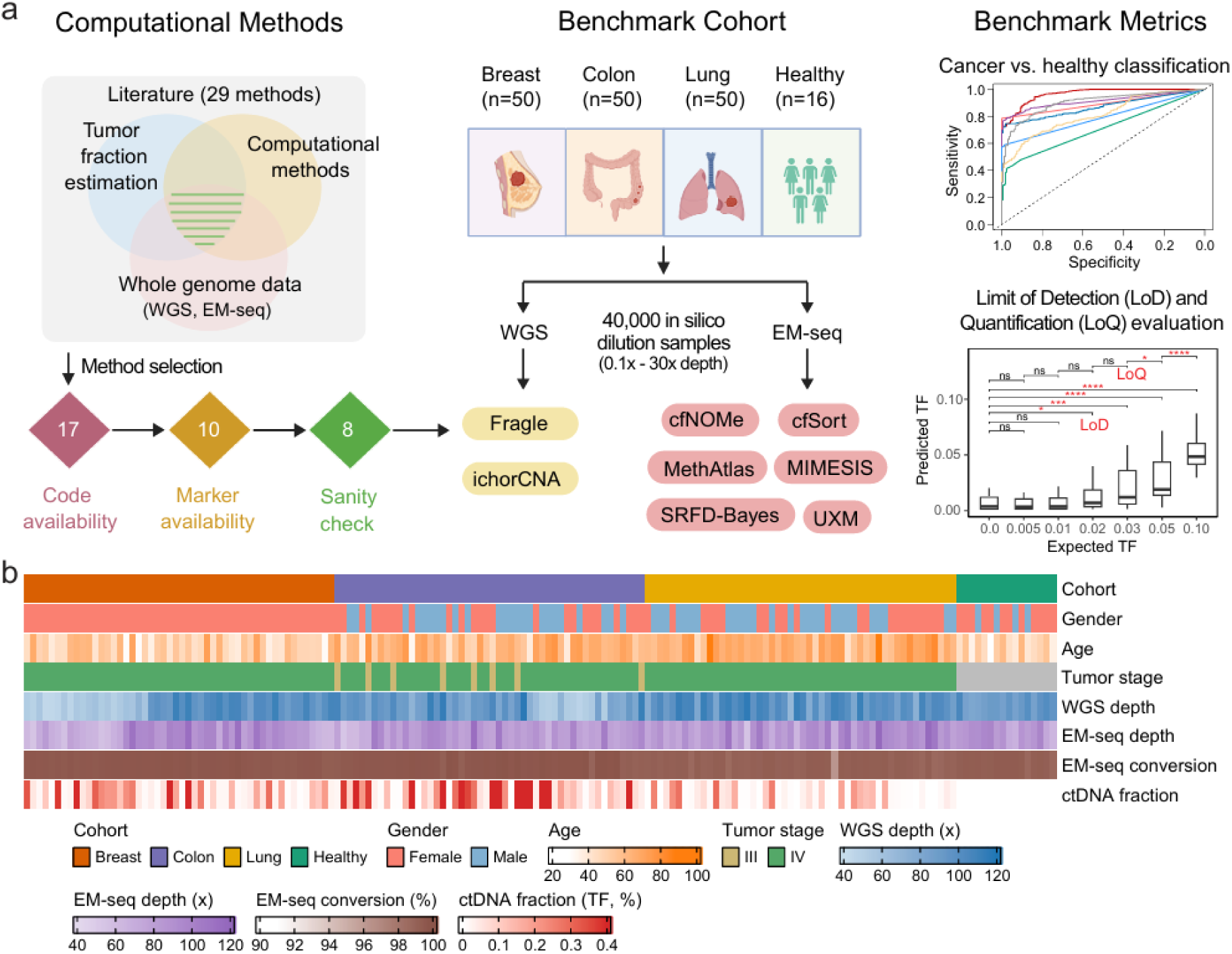
Overview of benchmarking approach. **b**) An exhaustive literature survey identified 29 candidate tumor-naïve computational methods for ctDNA tumor fraction (TF) estimation from WGS and EM-seq data, which were filtered to eight executable methods for benchmarking. Plasma cfDNA was collected from 150 patients with colorectal, lung or breast cancer and from 16 healthy individuals, followed by deep WGS and EM-seq profiling. Down-sampled technical replicates and in silico cancer-healthy dilution series were generated across sequencing depths and TFs. These datasets were used to assess detection accuracy, quantitative concordance, and limits of detection and quantification (LoD and LoQ). **b**) Overview of patient cohorts and sample characteristics.

### Evaluation of cancer versus healthy classification accuracy

We first assessed each method’s accuracy in distinguishing cancer from healthy samples across the three tumor types. Starting from high-depth original samples, we generated 10 down-sampled technical replicates per sample at fixed depths of 0.1x, 1x, 5x, and 30x (healthy only). The methods were applied to all down-sampled healthy and cancer samples. As a sanity check, all methods predicted significantly higher TF values in cancer samples than in healthy samples (Suppl. Fig. S2). We then evaluated the classification accuracy across cancer types and sequencing depths using the area under the curve (AUC), obtained by varying the tumor fraction (TF) threshold for each method. At an intermediate sequencing depth of 5x, WGS-based methods consistently outperformed methylation-based approaches across tumor types (Fig. 2a, Suppl. Fig. S3 for other sequencing depths). Fragle achieved the highest overall accuracy across all cancer types (mean AUC = 0.90), while ichorCNA (0.82) demonstrated best performance at high specificity (>95%) in breast and colorectal cancers. Among methylation-based methods at 5x depth, UXM showed the highest overall accuracy (mean AUC = 0.81), particularly in colorectal and breast cancers, where it was comparable to ichorCNA. Most methylation-based methods showed high accuracy at 30x, but accuracy declined at depths below 5x (Fig. 2b), with UXM and MethAtlas showing the least performance degradation at 0.1x. Among WGS-based methods, Fragle remained robust down to 1x, whereas ichorCNA retained accuracy even at 0.1x.

**Figure 2:**
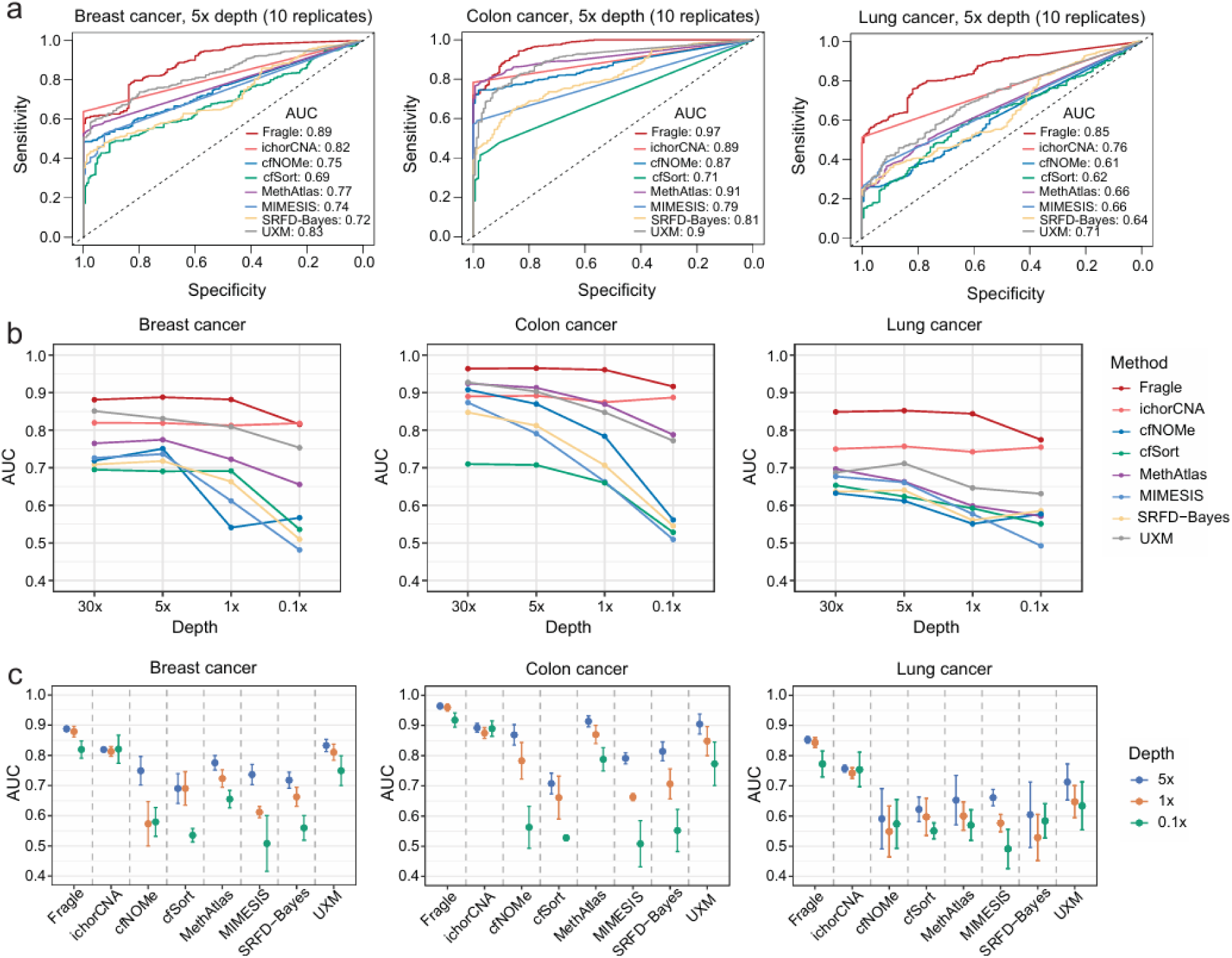
Cancer-healthy classification accuracy. **a**) Cancer–healthy classification accuracy (AUC) at 5x sequencing depth across three tumor types, based on 10 down-sampled technical replicates per sample. **b**) Classification accuracy as a function of sequencing depth (30x, 5x, 1x, and 0.1x). **c**) Variance in detection accuracy across down-sampled technical replicates. Points indicate the mean, and bars represent the standard deviation.

To further assess the impact of TF levels on classification performance, we stratified cancer samples into high- and low-TF groups (<10%). As expected, classification accuracy was highest in the high-TF samples and markedly lower in low-TF samples across all methods, with Fragle demonstrating superior accuracy in low-TF samples across all cancer types (Suppl. Fig. S4). By generating technical down-sampled replicates for each original sample, we estimated the variance in detection accuracy for each method (Fig. 2c). As expected, variance increased at lower sequencing depths, reflecting the greater impact of under-sampling and stochastic noise. ichorCNA also showed substantially higher variance at 0.1x, indicating that all tested methods benefit from at least 1x coverage for robust cancer detection. Overall, classification accuracy and robustness varied by method, tumor type, and sequencing depth, with all approaches showing increased variability at very low coverage and most methods requiring at least 1x depth for reliable classification.

### ctDNA quantification concordance

We next evaluated the quantitative characteristics of the methods across cancer types and sequencing depths. We first assessed the pairwise concordance of estimated TF values between methods. Most methods showed high inter-method correlation across cancer types at 5x sequencing depth (Fig. 3a, Suppl. Fig. S5 for other sequencing depths). In breast cancer samples, all method pairs exhibited strong correlation (Pearson *r* > 0.7). In colorectal cancer, only cfSort showed weak correlation with other methods (*r* < 0.4). In lung cancer samples, the correlation structure separated into two groups: one comprising both genomic and methylation-based methods (Fragle, ichorCNA, SRFD-Bayes and MIMESIS), and a second comprising only methylation-based methods (cfSort, UXM, MethAtlas and cfNOMe), potentially reflecting differences in the underlying methylation marker sets (Suppl. Fig. S6). Among WGS-based methods, ichorCNA tended to show higher correlation with methylation-based approaches, especially in breast and colon cancer samples (Fig. 3a).

**Figure 3:**
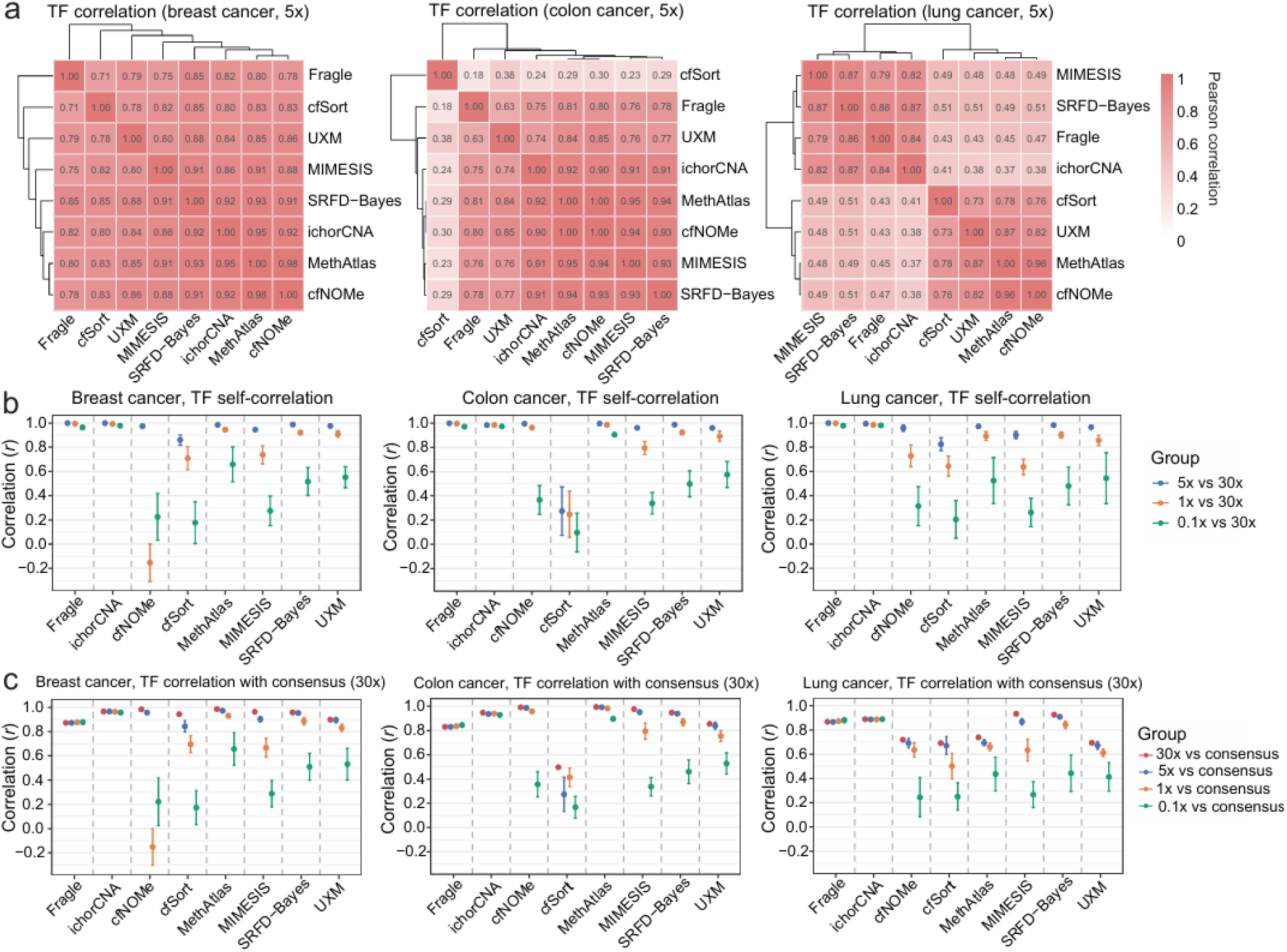
Quantification concordance. **a)** Pairwise Pearson correlation of TF estimates between methods at 5x sequencing depth for breast, colorectal, and lung cancer samples. **b)** Within-method concordance of TF estimates across sequencing depths (5x, 1x, 0.1x) relative to TF estimates at 30x depth. **c)** Correlation of individual methods with a consensus TF, defined as the median TF across all methods (30x depth), across cancer types and sequencing depths. Points represent means, with bars showing standard deviations.

To further examine the impact of sequencing depth on quantitative performance, we compared TF values obtained at the highest depth (30x) with those obtained at lower depths (5x, 1x, and 0.1x). Both ichorCNA and Fragle demonstrated high within-method concordance across all sequencing depths (Pearson’s *r* > 0.95; Fig. 3b). Most methylation-based approaches retained high concordance (*r* > 0.90) at 5x depth, but their robustness varied at lower depths. MethAtlas, SRFD-Bayes, and UXM had highest within-method concordance at 1x (*r* > 0.85) and 0.1x (*r* > 0.45). We next assessed the concordance of individual methods with a consensus TF, defined as the median across methods. IchorCNA showed high and consistent correlation with the consensus TF across all cancer types and sequencing depths (*r* > 0.87), whereas Fragle showed robust but slightly lower correlation (*r* > 0.82), particularly in breast and colorectal cancer (Fig. 3c). Among methylation-based methods, SRFD-Bayes showed the highest correlation across all cancer types, remaining robust down to 1x sequencing depth (*r* > 0.83). MIMESIS also maintained high correlation across cancer types at 5x depth (*r* > 0.86), but showed markedly reduced concordance at lower sequencing depths. Collectively, these results characterize how concordance among TF estimation methods varies by cancer type and sequencing depth.

### Limit of detection and quantification

To evaluate each method’s lower limit of detection (LoD) and quantification (LoQ), we generated a comprehensive sample dilution series. Consensus TFs were first estimated for all original samples as the median across all methods using 30x depth data. To ensure a fair benchmark, we selected samples from each cancer type with high and concordant ctDNA fractions across methods. For breast and colon cancer, we selected 10 samples with the lowest variance (median absolute deviation) and high median TF (>0.2). For the lung cancer cohort, where ctDNA fractions were lower, we selected 7 samples with the lowest variance and high TF (>0.15). Six healthy samples consistently predicted as low TF by all methods were selected for dilutions. Each cancer sample was stepwise diluted down to 0.5% TF, generating 30 technical replicates per dilution point using repeated subsampling and distinct healthy samples (Fig. 4a). The entire procedure was repeated at 30x, 5x, 1x, and 0.1x coverage to assess the impact of sequencing depth on LoD and LoQ. In total, this produced 38,880 in silico benchmark samples across the two data modalities.

**Figure 4:**
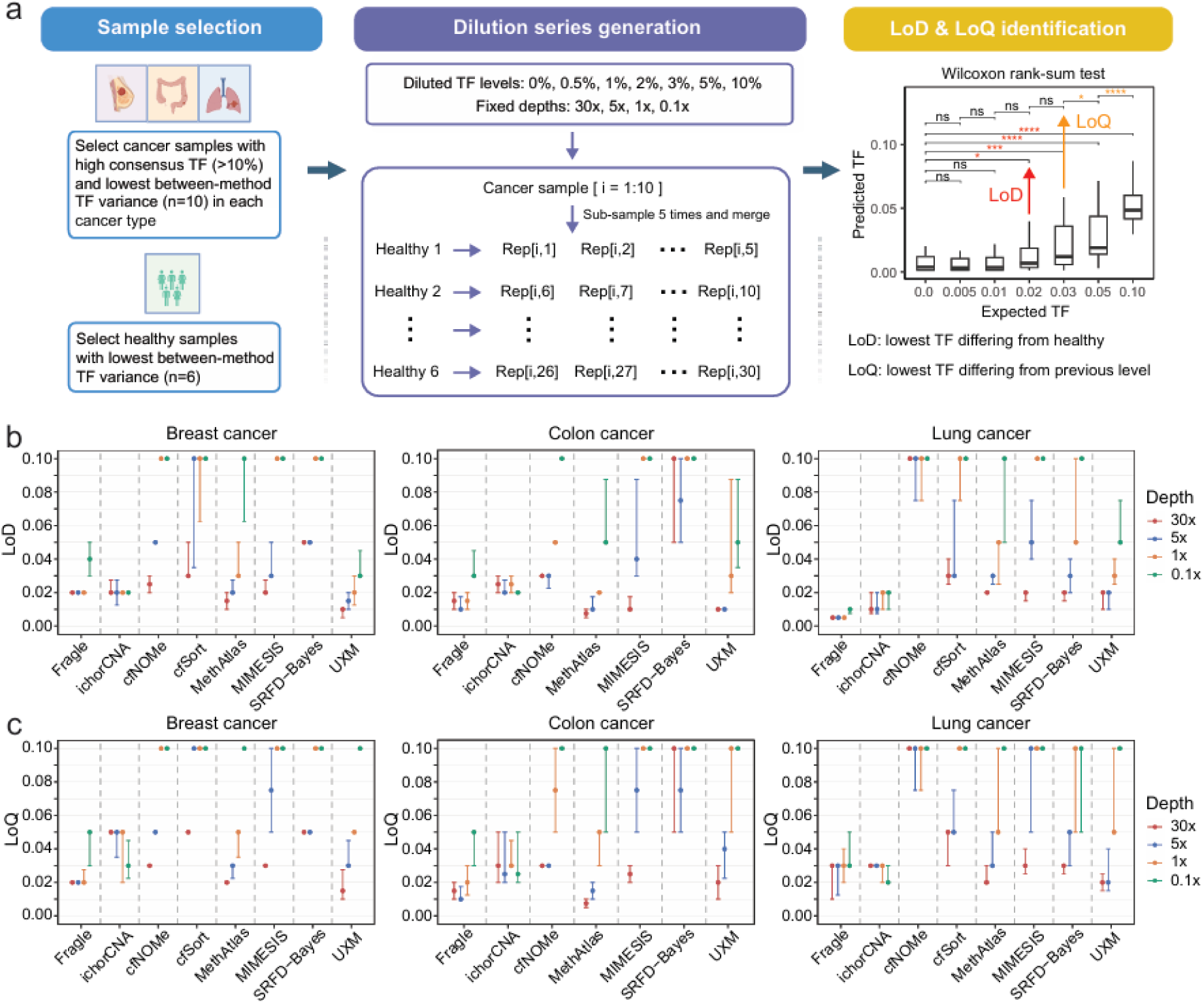
Limits of detection and quantification across methods and sequencing depths. **a**) Overview of the in silico dilution series generated from breast, colon, and lung cancer samples and healthy controls, with 30 replicates per dilution point, evaluated at 30x, 5x, 1x, and 0.1x coverage. **b, c**) Limit of detection (LoD) and quantification (LoQ) estimated for each method, cancer type, and sequencing depth. Points indicate medians across samples, with error bars showing inter-quartile ranges (25% and 75%).

To evaluate the LoD, we identified the lowest dilution point at which a method could distinguish cancer from healthy samples (Wilcoxon rank-sum test; Methods). The LoQ was defined as the lowest dilution point at which the method could distinguish that level from the next lower dilution level. In breast cancer, UXM achieved the lowest LoD at 30x (1%) and 5x (1.5%) coverage. Three other methods showed comparable LoD at 5x coverage: MethAtlas (2%), Fragle (2%), and ichorCNA (2%) (Fig 4b). At the lowest 0.1x coverage, ichorCNA outperformed other methods (2% vs. 3%). In colon cancer, MethAtlas had the lowest LoD (0.75% at 30x), followed by UXM (1% at 30x and 5x), MIMESIS (1% at 30x), and Fragle (1% at 5x). Fragle maintained a low LoD at 1x depth (1.5%), while the other methods showed a larger increase. In lung cancer samples, Fragle achieved the lowest LoD (0.5% at 1-30x) followed by ichorCNA (1% at 5-30x). Four methylation-based methods achieved a 2% LoD at 30x, with markedly worse performance at low coverage (0.1-1x).

We next evaluated the limit of quantification (LoQ) across methods. In breast cancer, UXM achieved the lowest LoQ at 30x depth (1.5%). At 5x depth, Fragle showed the lowest LoQ (2%), followed by MethAtlas and UXM (both 3%) (Fig. 4c). At 1x depth, Fragle maintained a low LoQ (2%), whereas MethAtlas and UXM exhibited a marked increase (5%). Notably, ichorCNA achieved its lowest LoQ (3%) at 0.1x depth, potentially reflecting its optimization for shallow WGS (<1x) data and suggesting that samples could be profiled or down-sampled to 0.1x depth to provide optimal LoQ performance. In colon cancer, MethAtlas (0.75% at 30x) achieved the lowest LoQ, followed by Fragle (1% at 5x). At 1x coverage, Fragle maintained the lowest LoQ (2%), whereas ichorCNA showed the best LoQ (3%) at 0.1x coverage. In lung cancer, IchorCNA (0.1x), UXM (5x-30x), and MethAtlas (30x) all achieved the lowest LoQ of 2%. Similar to other cancer types, only ichorCNA and Fragle maintained low LoQ (2–3%) at the lowest sequencing depths (0.1–1x).

In summary, methylation-based methods were more sensitive to sequencing depth for both LoD and LoQ, while WGS-based approaches showed relatively stable performance even at low coverage. Several methylation-based methods benefited substantially from high-coverage data, suggesting the potential for further performance gains at sequencing depths beyond 30x. However, limits of detection and quantification varied by cancer type for methylation-based methods, indicating that these approaches may also be constrained by challenges in selecting methylation marker sets specific to each cancer type.

### Assessment of multi-method and multi-modal approaches

Multi-method ensemble approaches can outperform individual methods when the constituent methods exhibit uncorrelated or complementary error profiles. We therefore systematically compared individual methods and their multi-method ensembles across sequencing depths across three tasks: Cancer-healthy classification, LoD, and LoQ. Double- and triple-method ensembles were generated using median TF estimates from the constituent methods. All approaches (single, double, and triple; *N* = 92) were then ranked by their average performance across the three cancer types. For cancer versus healthy sample classification, ensembles combining two WGS-based methods (Fragle and ichorCNA) showed consistently superior and robust performance across sequencing depths (Fig. 5a). While Fragle often showed strong performance as a single method, its combination with ichorCNA consistently yielded improved performance. Although both methods leverage WGS data, they interrogate distinct features – CNAs and fragment length profiles -likely providing complementary error profiles. Notably, methods based exclusively on methylation data were not ranked in the top 20 across any sequencing depth for cancer versus healthy sample classification (Suppl. Fig. S8).

**Figure 5:**
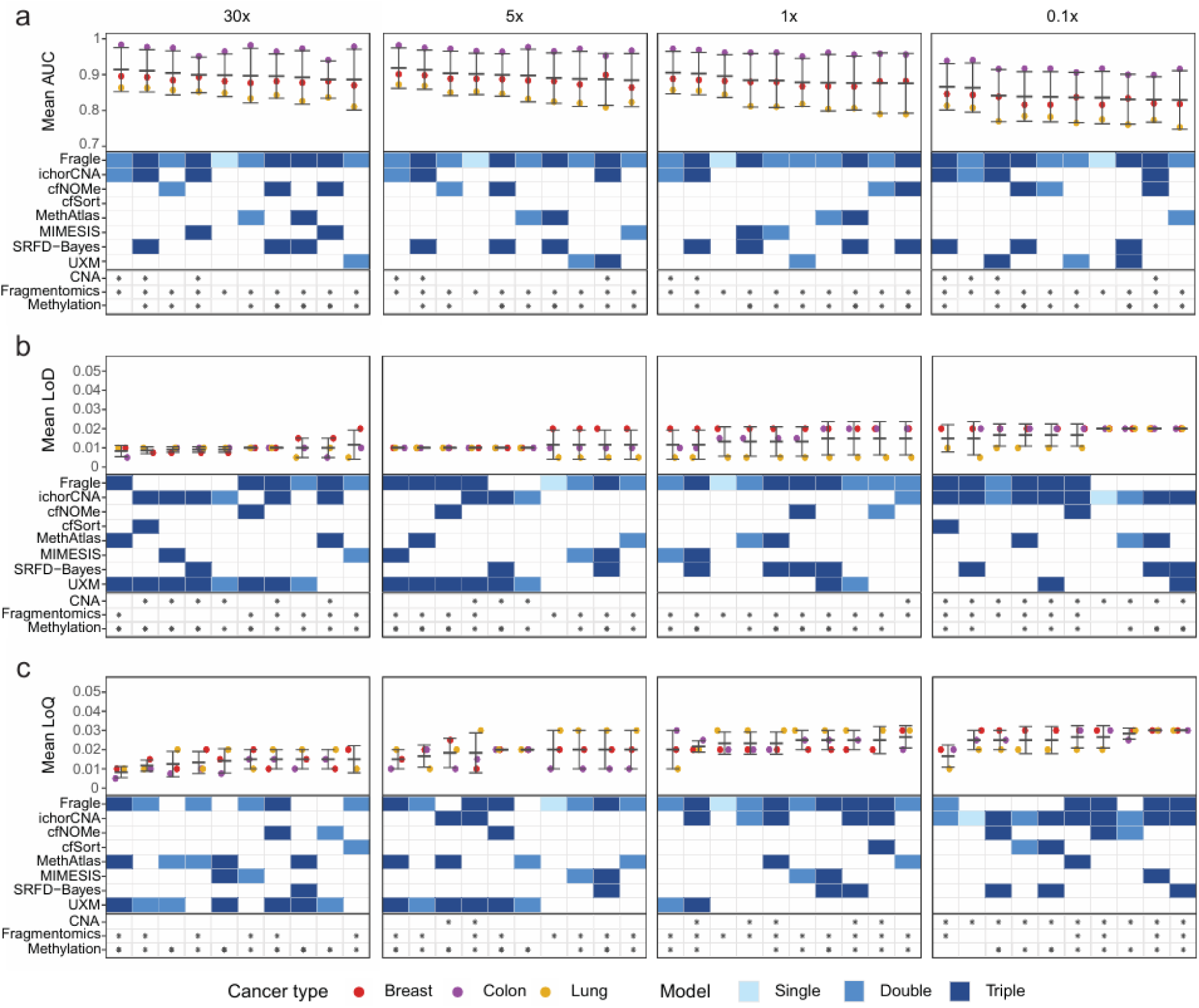
Multi-method performance evaluation across sequencing depths. **a**) Top-10 ranking of single-, double-, and triple-method ensembles (N = 92) for cancer versus healthy sample classification across distinct sequencing depths. Points represent the AUC for each cancer type, with bars indicating the mean and standard deviation. The feature modalities (CNA, Fragmentomics, Methylation) and method plurality (single, double, and triple) are indicated for each ensemble. **b**) Ranking of ensemble models based on their limit of detection (LoD) estimated from the sample dilution series. Points show the median LoD for each cancer type, with bars indicating the mean and standard deviation. cfSort was excluded in LoD/LoQ analysis for colon cancer due to its low TF correlation with all other methods. **c**) Ranking of ensemble models based on their limit of quantification (LoQ) estimated from the sample dilution series. Points show the median LoQ for each cancer type, with bars indicating the mean and standard deviation.

We next evaluated whether multi-method ensemble approaches could improve the limit of detection (LoD) and limit of quantification (LoQ). To this end, we applied all 92 ensemble combinations to the previously generated dilution series (Fig. 4a). At the highest sequencing depth (30x), method combinations involving both methylation and WGS-based approaches showed superior performance, achieving an LoD of approximately 1% (Fig. 5b). The lowest average LoD across the three cancer types was observed for a combination of three methods (Fragle, MethAtlas, UXM), although other combinations pairing UXM with either of the two WGS-based methods also performed well. At 5x and 1x coverage, most top-performing combinations again integrated both WGS- and methylation-based methods, with Fragle being the only single-method approach showing comparable performance. At the lowest sequencing depth (0.1x), combinations incorporating both WGS-based methods generally performed well, with ichorCNA being the only single-method approach that showed a robust LoD.

Evaluation of LoQ at 30x coverage showed a similar pattern, with all top-10 combinations including at least one methylation-based method and combinations involving UXM and MethAtlas performing best (Fig. 5c). At 5x and 1x coverage, combinations involving both WGS and methylation methods showed superior LoQ. Fragle was the only single-method approach represented in the top 10 at these coverage levels. At the lowest sequencing depth (0.1x), the combination of Fragle and ichorCNA showed the best performance, with ichorCNA also demonstrating robust single-method performance. Overall, these results highlight that integrating complementary WGS and methylation features often enhances detection sensitivity and quantification across cancer types. In our benchmark, the optimal combination of methods depends on sequencing depth and whether the focus is on classification, detection, or quantification.

### Prioritisation of data modalities and sequencing depth across cancer types

Practical implementation of multi-modal ensemble approaches often involves a cost-benefit trade-off, as higher sequencing depth and multiple data modalities increase cost. We therefore performed a focused analysis within individual cancer types, ranking ensemble methods across sequencing depths and data modalities. For each cancer type, we selected the top-performing ensemble method for every combination of sequencing depth (0.1x–30x) and data modality (methylation, WGS, or both). When multiple methods were tied, we prioritized simpler ensembles (lower plurality) or included all tied methods. In cancer versus healthy classification, WGS-based combinations outperformed methylation-based approaches (Fig. 6a). For multimodal approaches, the incorporation of methylation data yielded only a modest performance gain over WGS alone at 30x coverage in breast cancer and did not improve performance at lower coverage levels or in other cancer types. When models were ranked by average performance across all three cancer types (pan-cancer), WGS-based methods also showed the highest accuracy (Fig. 6b). Their performance remained stable from 1x to 30x coverage (AUC 0.91–0.92), with a decline at 0.1x coverage (AUC 0.86). Multimodal approaches did not outperform WGS-only methods, and methylation-only approaches – even at 30x coverage – showed substantially lower classification accuracy (AUC 0.82).

**Figure 6:**
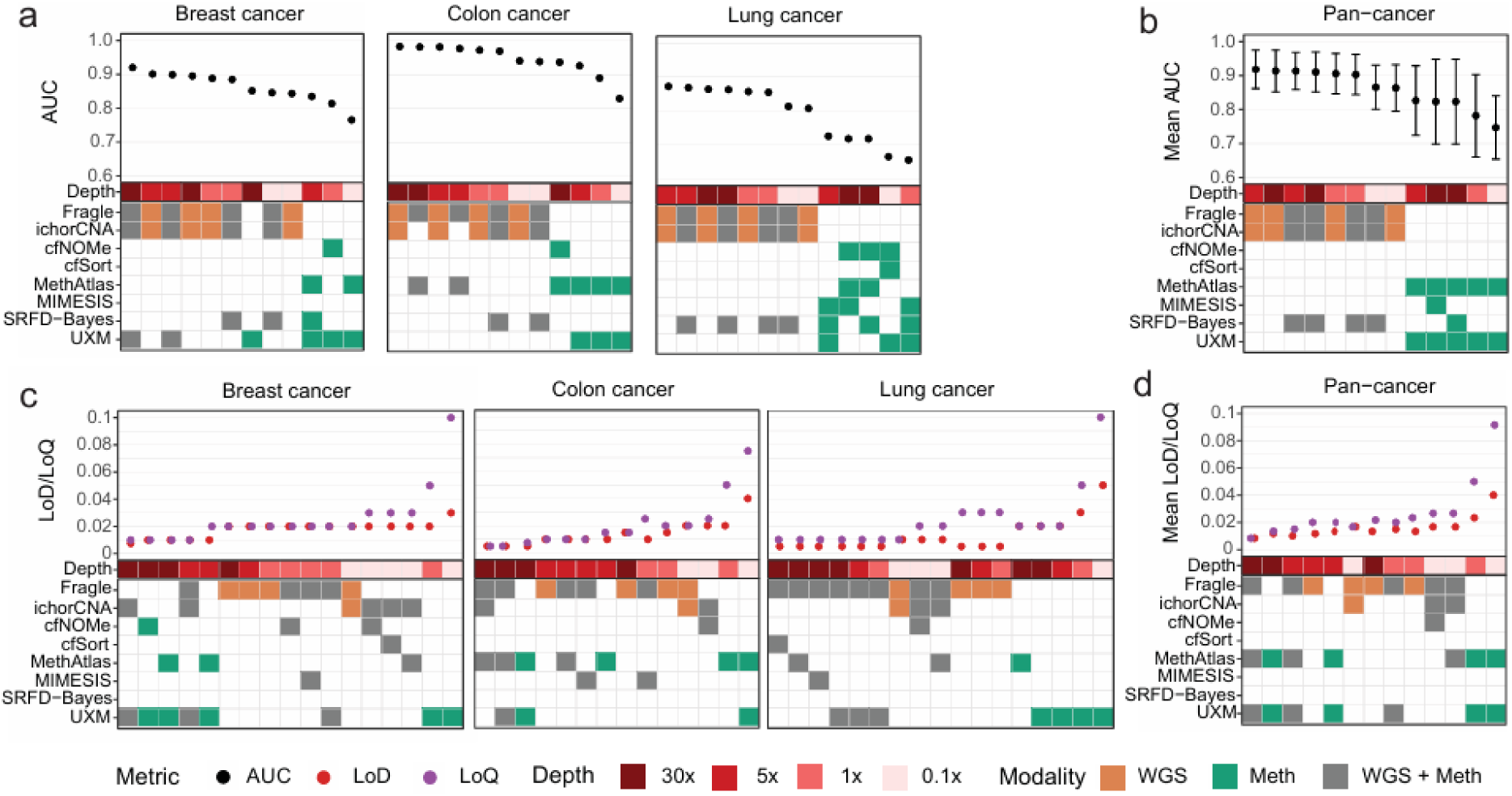
Prioritization of data modalities and sequencing depth across cancer types. **a**) Top-performing ensemble methods for each combination of sequencing depth (0.1x–30x) and data modality (methylation, WGS, or both) for cancer versus healthy classification (AUC). Ties were resolved by prioritizing simpler ensembles (lower plurality) or including all tied methods. **b**) Same analysis, with ensemble methods ranked by pan-cancer performance (average AUC across the three cancer types). **c**) Top-performing ensemble methods for limits of detection (LoD) and quantification (LoQ). **d**) Same analysis, with ensemble methods ranked by pan-cancer performance (average LoD and LoQ across the three cancer types).

A similar analysis of the limit of detection (LoD) and quantification (LoQ) across method combinations revealed a more complex decision landscape. Approaches combining methylation and WGS, closely followed by methylation-only approaches, generally achieved superior detection and quantification limits at the highest coverage (30x; Fig. 6c). However, WGS-only approaches at lower coverage often yielded comparable or only slightly reduced performance. In breast cancer, a 1x WGS-only approach (2% LoD/LoQ) achieved moderately reduced performance compared to the best 30x multi-modal approach (0.75% LoD, 1% LoQ). In colon cancer, a 5x WGS-only approach (1% LoD/LoQ) showed slightly lower performance than the best 30x multi-modal approach (0.5% LoD/LoQ). In lung cancer, a 0.1x WGS-only approach (1% LoD/LoQ) achieved performance approaching the best 30x multi-modal approach (0.5% LoD, 1% LoQ). When models were ranked by average performance across all three cancer types, a 30x multi-modal approach achieved the best limits of detection and quantification (∼1% LoD/LoQ). However, the performance of 30x methylation-only (LoD 1.2%, LoQ 1.3%) and 5x WGS-only (LoD 1.2%, LoQ 2%) approaches were only marginally lower. Collectively, these analyses define a practical framework for selecting sequencing modality and depth, highlighting that the marginal benefits of high coverage approaches must be weighed against cost and cancer-type dependent performance characteristics.

## Discussion

Circulating tumor DNA analysis is often pursued using tumor-naïve, genome-wide approaches such as low-pass whole-genome sequencing (lpWGS) and DNA methylation profiling. From an initial set of 29 published computational methods, we identified eight methods with available software for ctDNA quantification based on distinct genome-wide features – copy number alterations, fragmentomics, and methylation. Using matched deep WGS and EM-seq data from 150 cancer plasma samples, we generated >40,000 in silico samples to provide a comprehensive assessment of method performance across sequencing depths, tumor fractions, and cancer types.

Our results indicate that two WGS-based methods, ichorCNA and Fragle, show the strongest performance at low sequencing coverage among the approaches evaluated, with further improvements observed when the methods are combined. Methylation-based methods often achieved superior limits of detection and quantification at high coverage (30x), however, their incremental benefit over lower-coverage WGS approaches was often context-dependent. Multi-modal ensemble approaches that integrate WGS and methylation data can further improve performance at high sequencing coverage, but the optimal combination is dependent on sequencing depth, cancer type, and whether the focus is on classification, detection, or quantification. Interestingly, our analysis revealed a divergence in the performance of methylation-based methods for cancer–healthy classification and LoD/LoQ tasks. Methylation-based methods showed excellent LoD/LoQ at high sequencing coverage but notably reduced cancer–healthy classification performance. Moreover, the methods that performed best in LoD/LoQ tasks (UXM, MethAtlas) differed from those that best complemented WGS-based approaches in classification (SRFD-Bayes). This divergence suggests that methylation-based approaches may be more adversely affected by inter-tumor heterogeneity than WGS-based methods.

Our results also highlight opportunities for method development and expanded benchmarking. Only methylation-based approaches benefited from sequencing depths above 5x, indicating potential to optimize WGS-based methods at higher coverage. Several methylation-based methods showed notable performance gains with high- coverage data, indicating that sequencing depths beyond 30x may yield further improvements, in line with previous studies of high-coverage targeted methylation assays (27). This study focused on ctDNA quantification and did not evaluate other whole-genome methods developed solely for cancer detection, which represents an important avenue for future work.

Taken together, this benchmark provides a framework for selecting fit-for-purpose, tumor-naïve whole-genome strategies for ctDNA quantification. For applications prioritizing robust detection and quantification across multiple tumor types, low- to mid-depth WGS profiling represents an efficient approach, whereas high-coverage multimodal strategies may be advantageous when higher-resolution quantification is required. Identification of fit-for-purpose ctDNA profiling approaches is increasingly important as the field explores ctDNA kinetics as a quantitative readout of therapeutic response across distinct treatment modalities and cancer types (8–12). By establishing the optimal conditions for scalable single- and multi-modal ctDNA profiling, our benchmark provides a rigorous foundation to accelerate methodological development and support new ctDNA research studies.

## Methods

### Patient cohort

Volunteers were recruited at the National Cancer Centre Singapore under studies 2018/2795 (colorectal cancer), 2018/2963 (lung cancer), 2018/2709 (breast cancer), and 2019/2401 (healthy volunteers). All studies were approved by the SingHealth Centralised Institutional Review Board, and written informed consent was obtained from all participants. Clinical data for the included patients are provided in Suppl. Table S2.

### Plasma samples

Plasma was separated from blood within 2 hours of venipuncture by centrifugation at 300 x g for 10 minutes, followed by 9,730 x g for 10 minutes, and stored at −80 °C. Samples were shipped overnight on dry ice and stored at −80 °C until further processing. Cell-free DNA (cfDNA) was extracted from 4 mL of plasma using the KingFisher Flex sample purification kit (Thermo Fisher Scientific, Fremont, CA, USA) and quantified with the Qubit 1X dsDNA High Sensitivity Assay kit (Thermo Fisher Scientific). cfDNA quality and high molecular weight contamination were assessed by qPCR using a modified KAPA NGS FFPE DNA QC protocol with custom cfDNA-specific primers (Roche Sequencing Solutions, Pleasanton, CA, USA).

### WGS and EM-seq library preparation and sequencing

cfDNA libraries were prepared from 2 ng cfDNA using the KAPA EvoPrep Kit (Roche Sequencing Solutions, Pleasanton, CA, USA) with 0.2 uM KAPA Universal Adapters. Input mass was adjusted to account for high molecular weight contamination measured by qPCR. All libraries were amplified for 8 PCR cycles using the KAPA UDI Primer Mixes and KAPA HiFi HS Readymix (Roche Sequencing Solutions, Pleasanton, CA, USA). After PCR, libraries underwent a double 1x clean-up post PCR with KAPA HyperPure Beads to ensure removal of adapter dimers. Libraries were quantified using the Qubit 1X dsDNA High Sensitivity Assay kit (Thermo Fisher Scientific, Fremont, CA, USA) and Agilent High Sensitivity D1000 ScreenTape System (Agilent Technologies, Waldbronn, Germany). Enzymatic methyl-converted cfDNA libraries were prepared from 2 ng cfDNA using the KAPA EvoPrep Kit (Roche Sequencing Solutions, Pleasanton, CA, USA) with an initial treatment with a proprietary DNA enhancer, followed by processing with the NEBNext Enzymatic Methyl-seq Conversion Module (New England Biolabs, Ipswich, MA, USA) with 0.22 uM custom methylated adapters. Input mass was adjusted to account for high molecular weight contamination measured by qPCR. All libraries were sequenced on NovaSeq 6000 instruments (Illumina, San Diego, CA, USA) using paired-end 150 bp read length to achieve a mean whole-genome coverage of at least 40x. A pooling strategy was adopted to minimise any possible batch effects. Each sequencing pool contained a mixture of plasma samples from cancer patients and healthy individuals, and pools were sequenced across multiple S4 flow cells.

### Processing of WGS and EM-seq data

Adapter sequences were trimmed from raw paired-end whole-genome sequencing (WGS) and enzymatic methyl-sequencing (EM-seq) reads using Fastp (RRID:SCR_016962; version 0.20.0; cut_window_size=5; qualified_quality_phred=20; length_required=30) (28). After trimming, WGS reads were aligned to the human reference genome GRCh38 using BWA-MEM (RRID:SCR_010910; BWA v0.7) (29). EM-seq reads were aligned to GRCh38 and deduplicated using Bismark (RRID:SCR_005604; v0.24.2) (30), with Bowtie2 (RRID:SCR_016368; v2.5.4) (31) used as the internal aligner (default settings). For both WGS and EM-seq datasets, aligned reads were subjected to quality control using Samtools (RRID:SCR_002105; v1.15) (32). Sequencing depth was calculated using PanDepth (v2.25) (33). Both WGS and EM-seq BAM files were down-sampled to target coverage levels of 0.1x, 1x, 5x, and 30x using Samtools. DNA methylation calling for EM-seq data was performed using Bismark.

### ctDNA quantification methods

We surveyed 29 published computational methods for WGS- and methylation-based ctDNA tumor fraction estimation (published before January 2025). Eight methods – two WGS-based and six methylation-based – were selected based on publicly available software, required DNA methylation biomarker sets, and successful execution and validation on test data (Suppl. Table S1): (1) FRAGLE (16), (2) ichorCNA (13), (3) cfNOMe (22), (4) cfSort (23), (5) MethAtlas (1), (6) MIMESIS (24), (7) SRFD-Bayes (25), and (8) UXM (23). Methods lacking a usable software tool, pre-trained model, or tissue-specific methylation marker sets were excluded from the benchmark.

FRAGLE and ichorCNA were applied to WGS data, whereas the remaining six methods were applied to EM-seq data. For all methylation-based methods, tumor fraction estimation was performed using the cell-type/tissue-specific reference biomarker panels distributed with the respective software packages. All methods were executed using their default parameters. All methods were applied to reads aligned to the GRCh38 reference genome. cfSort provides markers only for hg19, so its input files were converted to GRCh37 coordinates using CrossMap (RRID:SCR_001173; v0.7.0). For UXM, PAT files required for downstream analysis were generated using wgbstools (https://github.com/nloyfer/wgbs_tools). UXM was implemented using the two sets of reference markers provided with the package, and the mean tumor fraction values were subsequently utilized for analysis.

### ROC analysis

Receiver operating characteristic (ROC) curves were generated using the pROC R package, with tumor fraction (TF) values estimated by each method used as the continuous predictor variable.

### Generation of in silico dilution series

Cancer plasma samples with high median TF (> 0.2) and low inter-method variability (median absolute deviation, MAD < 0.2) across the eight TF estimation methods were selected for generation of the in silico dilution series. For lung cancer, this minimum TF limit was set to 0.15 due to the lack of high TF samples. In total, 27 cancer samples (10 breast cancer, 10 colon cancer, and 7 lung cancer samples) were selected based on these 2 criteria. In addition, six healthy samples with the lowest variability (MAD) were selected as healthy controls for dilutions. For each selected cancer sample, five independent random subsamples of sequencing reads were extracted and mixed with randomly sampled reads from the six healthy samples to generate 30 dilution mixtures per cancer sample at each sequencing depth and at each TF level. The proportions of cancer-derived and healthy-derived reads were varied to generate dilution samples across a range of tumor fractions: 0%, 0.5%, 1%, 2%, 3%, 5%, and 10%. The total number of sampled reads was selected to simulate four sequencing coverage levels: 0.1x, 1x, 5x, and 30x. For the LoD/LoQ benchmark, we generated 38,880 in silico dilution cancer samples by combining 27 original cancer samples, 30 dilutions per sample, 6 tumor-fraction levels, 4 sequencing depths, and 2 data modalities (WGS and EM-seq). The 16 healthy samples were down-sampled 10 times at all four sequencing depths across both data modalities, yielding 1,280 healthy in silico replicates. Altogether, the benchmark included 40,160 in silico samples. Because sex-associated differentially methylated regions could theoretically introduce bias in mixed-sample dilutions, we evaluated the impact of sex matching. No differences were observed between sex-matched and sex-unmatched dilution series for any DNA methylation-based method, as determined by concordance between predicted and true tumor fractions (Suppl. Fig. S7).

### Limit of detection (LoD) and limit of quantification (LoQ) analysis

Limits of detection (LoD) and quantification (LoQ) for each tumor-fraction estimation method were determined using a nonparametric hypothesis-testing framework applied to the in silico dilution series. The LoD for a given method and sample was defined as the lowest tumor-fraction level at which the predicted TF distribution (N = 30 replicates) was significantly higher than the healthy background (N = 160 replicates), using a one-sided Wilcoxon rank-sum test with a significance threshold of α = 0.01. The LoQ was defined as the lowest tumor-fraction level (e.g., 2%) at which the predicted TF distribution was significantly higher than that of the immediately lower dilution level (e.g., 1%), again assessed using a one-sided Wilcoxon rank-sum test with α = 0.01.

### Ensemble models

We generated ensemble models by combining methods in all possible double- and triple-method configurations. For each ensemble, the median TF value across its constituent methods was calculated and used as the combined TF estimate for ROC, LoD, and LoQ analyses. cfSort was excluded in detection and LoD/LoQ analysis for colon cancer due to its low TF correlation with all other methods (*r* < 0.4; Fig. 3a).

### Code availability

All scripts, configuration files, and code required to reproduce the analyses and figures are publicly available on GitHub at https://github.com/skandlab/ctDNA-quant-benchmark.

## Acknowledgements

This work was supported by the Singapore Ministry of Health’s National Medical Research Council under its OF-IRG program (OFIRG21nov-0083), with support by the A*STAR Computational Resource Center through the use of its high performance computing facilities.

