## Supplemental figure S1-S8 for "Systematic benchmarking of multi-modal approaches for tumor-naïve ctDNA detection and quantification"

**Supplementary Document**

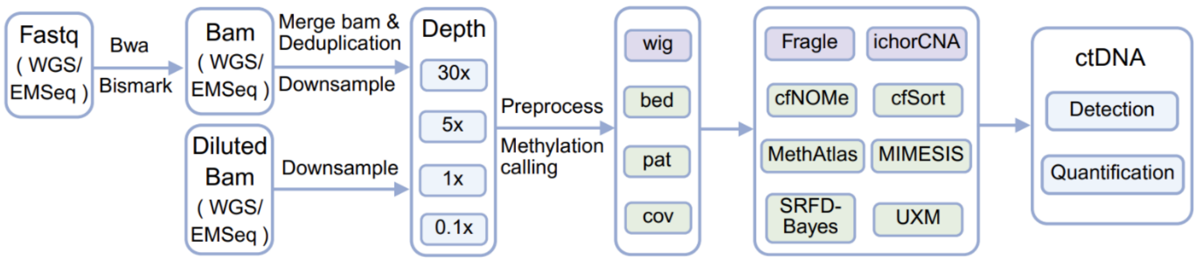


**Figure S1. Schematic workflow of the computational pipeline.** Trimmed FASTQ sequencing read-pairs from Whole Genome Sequencing (WGS) and Enzymatic Methyl-seq (EM-seq) are aligned using BWA and Bismark, respectively. Resulting BAM files are merged, deduplicated, and down sampled to multiple depths (30x, 5x, 1x, and 0.1x). Genomic and DNA methylation-level features are extracted in standardized formats (wig, bed, pat, and cov) as required for downstream analysis. These data are integrated into a multi-modal suite comprising eight computational tools benchmarked in this study for the detection and quantification of ctDNA.

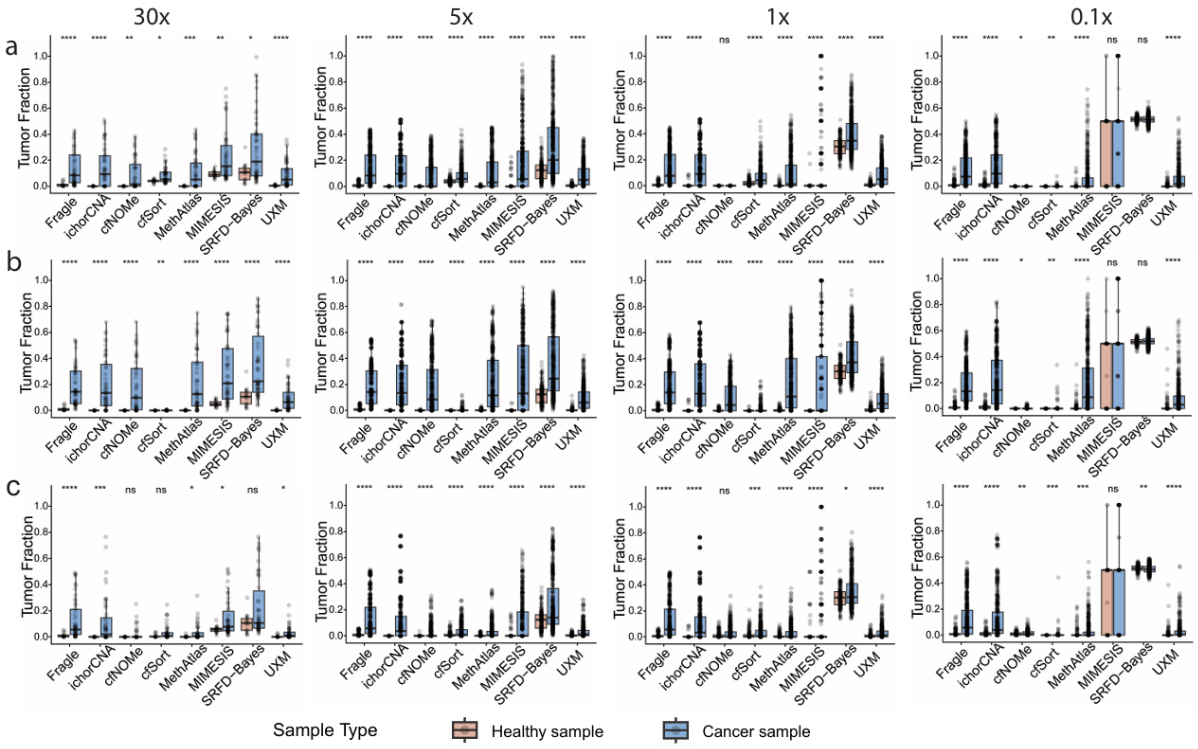


**Figure S2. Tumor fraction estimates in healthy and cancer samples.** The eight tumor fraction estimation methods were applied to plasma samples from healthy volunteers and patients with three cancer types across four sequencing depths: **a**) breast cancer, **b**) colon cancer, **c**) lung cancer. A Wilcoxon rank-sum test was used to determine if tumor fraction (TF) values differed significantly between healthy and cancer samples.

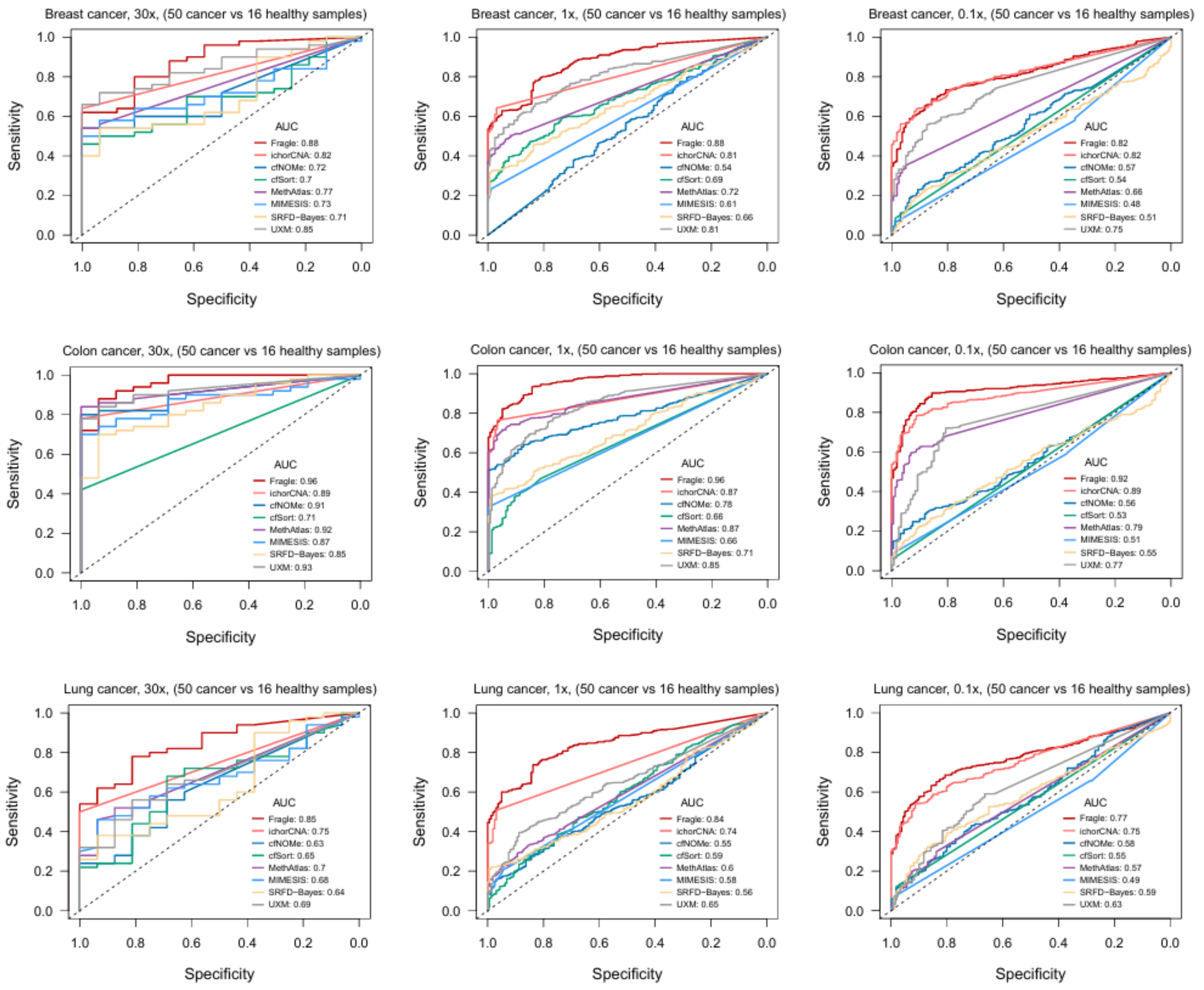


**Figure S3. Cancer vs. healthy classification accuracy across methods and sequencing depths.** Receiver operating characteristic (ROC) curves for cancer–healthy classification across the three tumor types (Breast, Colon, and Lung), three sequencing depths (30x, 1x, and 0.1x; 5x in main manuscript). Plots for 1x and 0.1x depths are based on 10 down-sampled technical replicates per sample, while the 30x data represents the original samples without technical replicates (number of samples / technical replicates indicated above each plot).

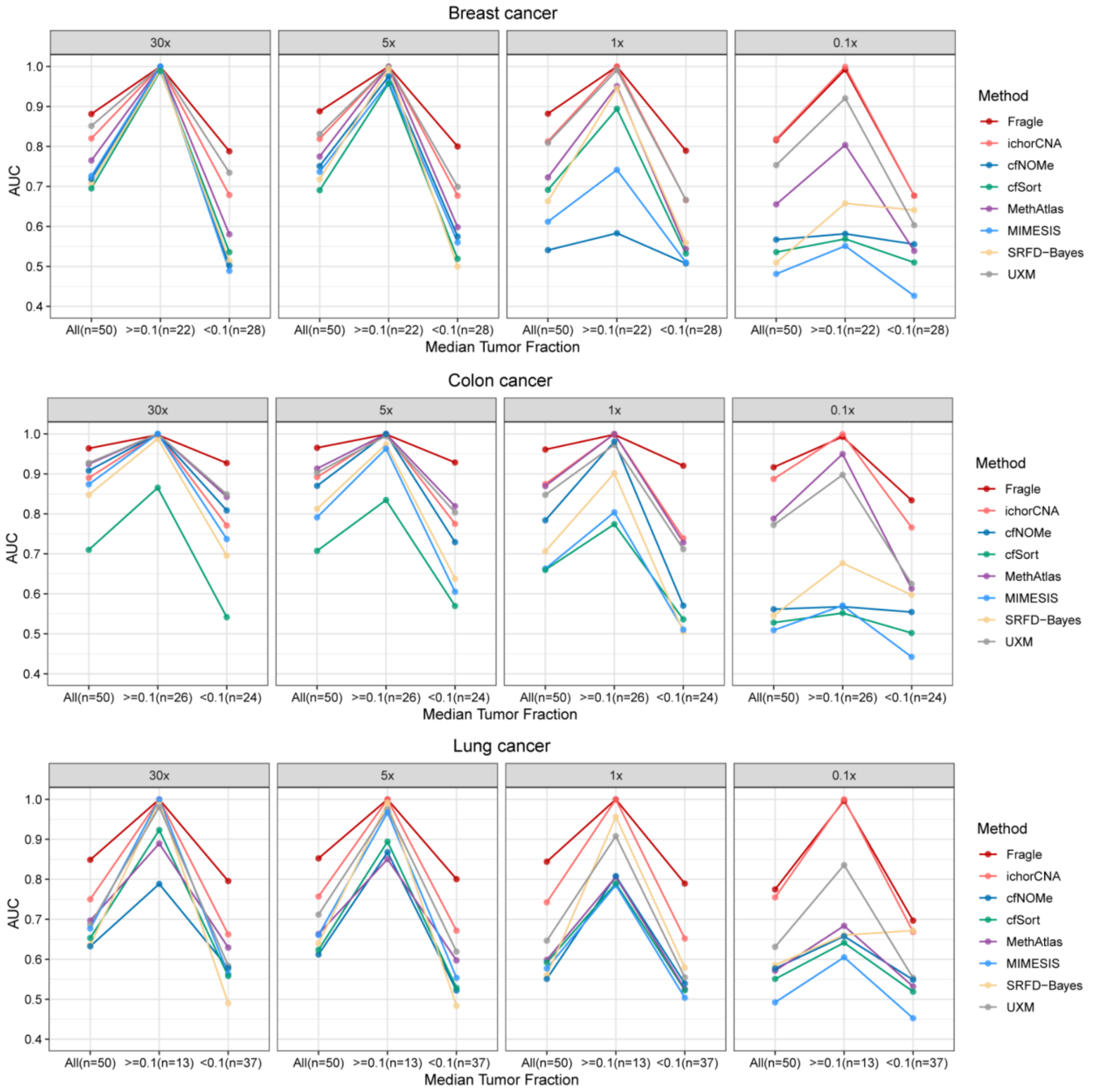


**Figure S4. Classification performance at high and low tumor fractions.** Area under the curve (AUC) values for cancer–healthy classification stratified by median tumor fraction (TF): all samples, high TF (≥ 0.1), and low TF (< 0.1). Results are displayed for eight estimation methods across three tumor types (Breast, Colon, and Lung) and four sequencing depths (30x, 5x, 1x, and 0.1x). Points represent the AUC for each specific subgroup.

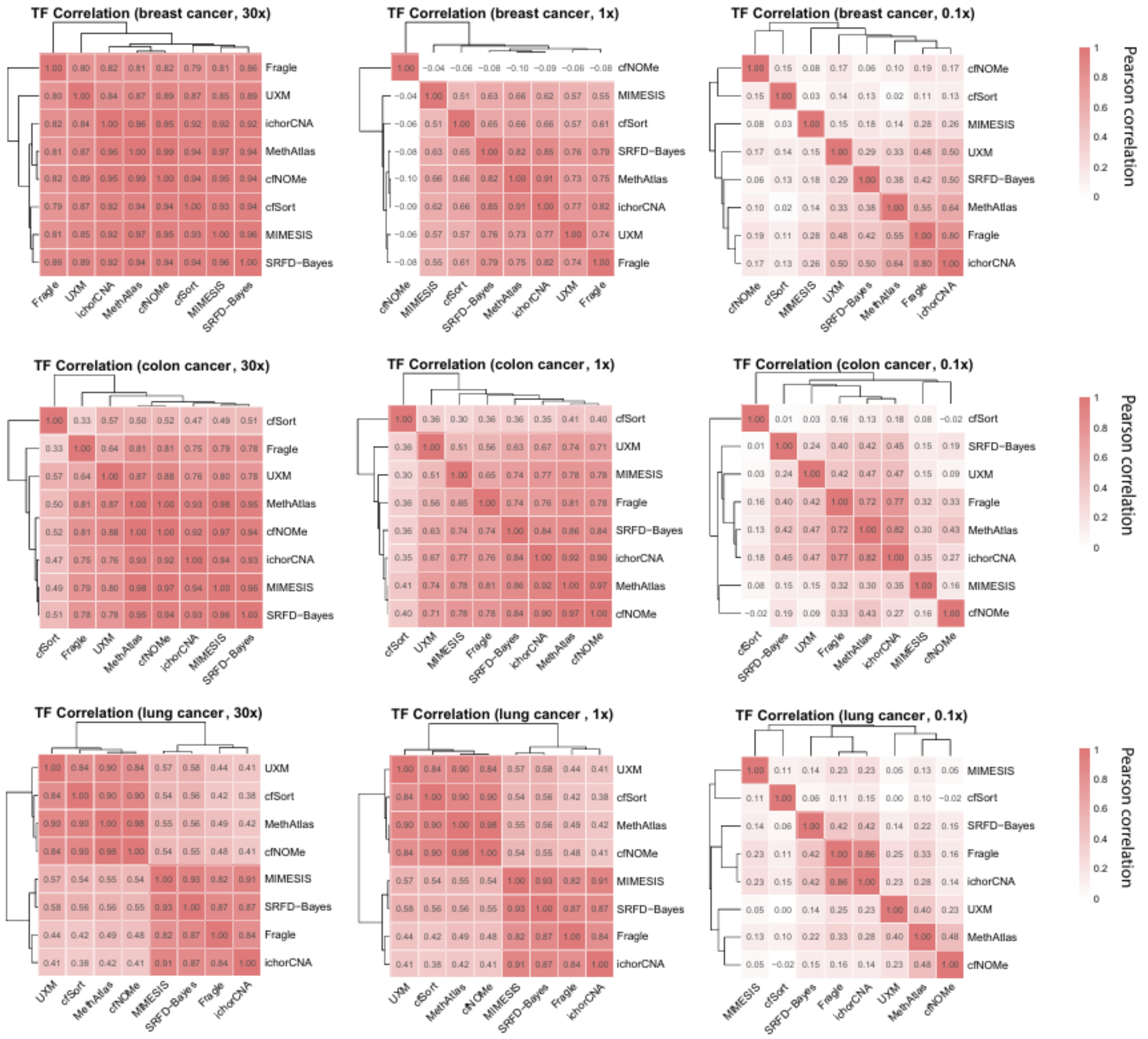


**Figure S5. Concordance of tumor fraction quantification across methods.** a) Heatmaps displaying pairwise Pearson correlation coefficients ($r$) of tumor fraction (TF) estimates across eight methods. Data are shown for three cancer types (Breast, Colon, and Lung) across four sequencing depths (30x, 1x, and 0.1x). The dendrograms indicate hierarchical clustering based on correlation concordance (method similarity).

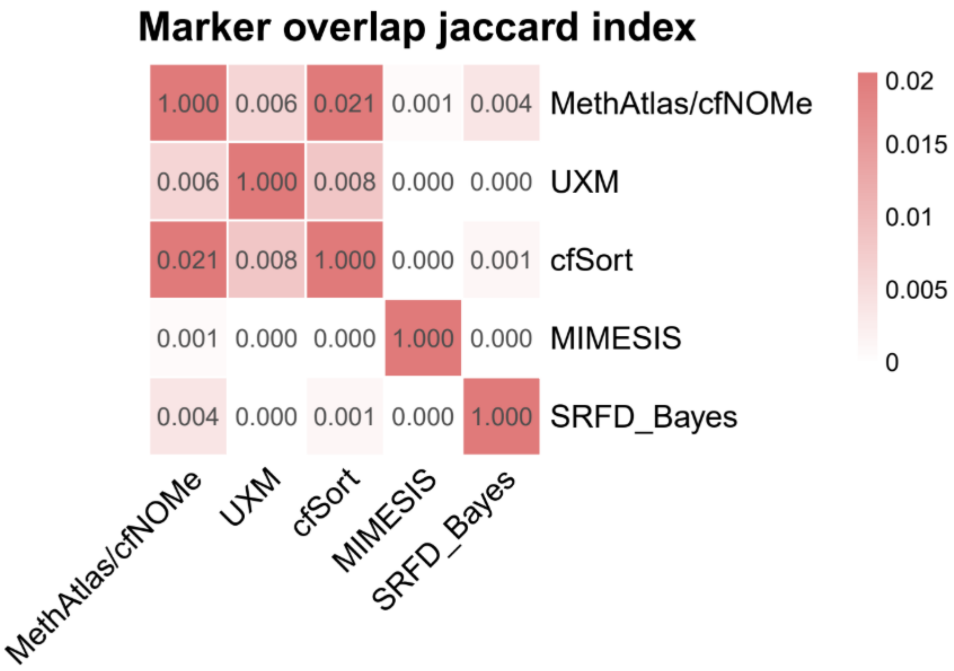


**Figure S6. Marker similarity between methylation-based deconvolution methods.** The heatmap quantifies the similarity between the reference tissue/cell-type methylation marker sets utilized by six distinct tumor fraction estimation tools. Values represent the Jaccard similarity index, calculated as the size of the intersection divided by the size of the union of the marker sets. The diagonal values represent the self-overlap of each tool's specific marker set.

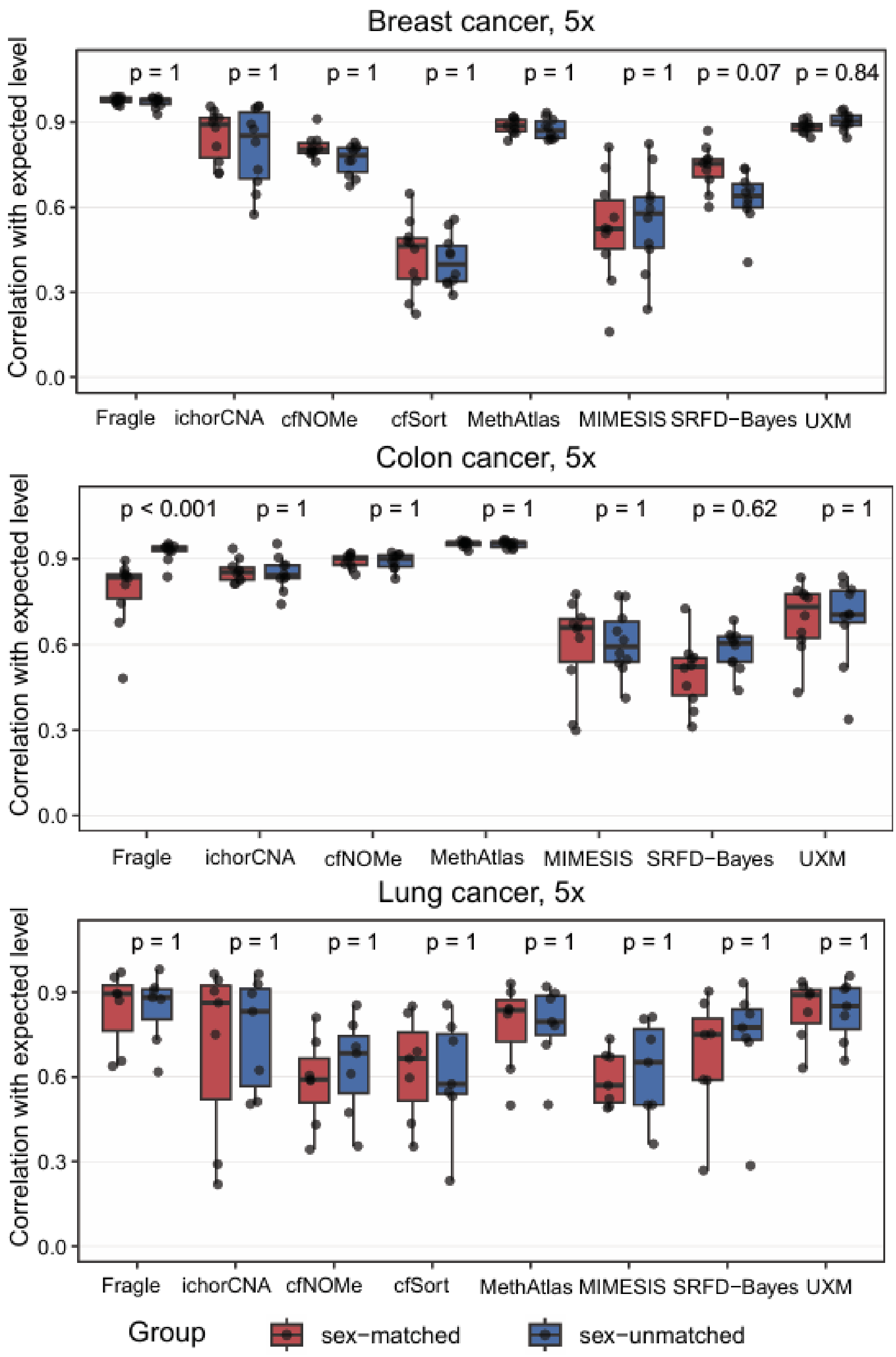


**Figure S7. Impact of sex-matched and sex-unmatched dilution samples**. Samples were diluted using sex-matched (red) and sex-unmatched (blue) dilution samples, correlation between predicted tumor fractions and diluted (expected) levels for breast, colon, and lung cancer samples at 5x depth. Points represent individual samples. Adjusted P-values (Bonferonni-corrected, Wilcoxon rank-sum test) indicate differences between sex-matched and sex-unmatched groups for each method.

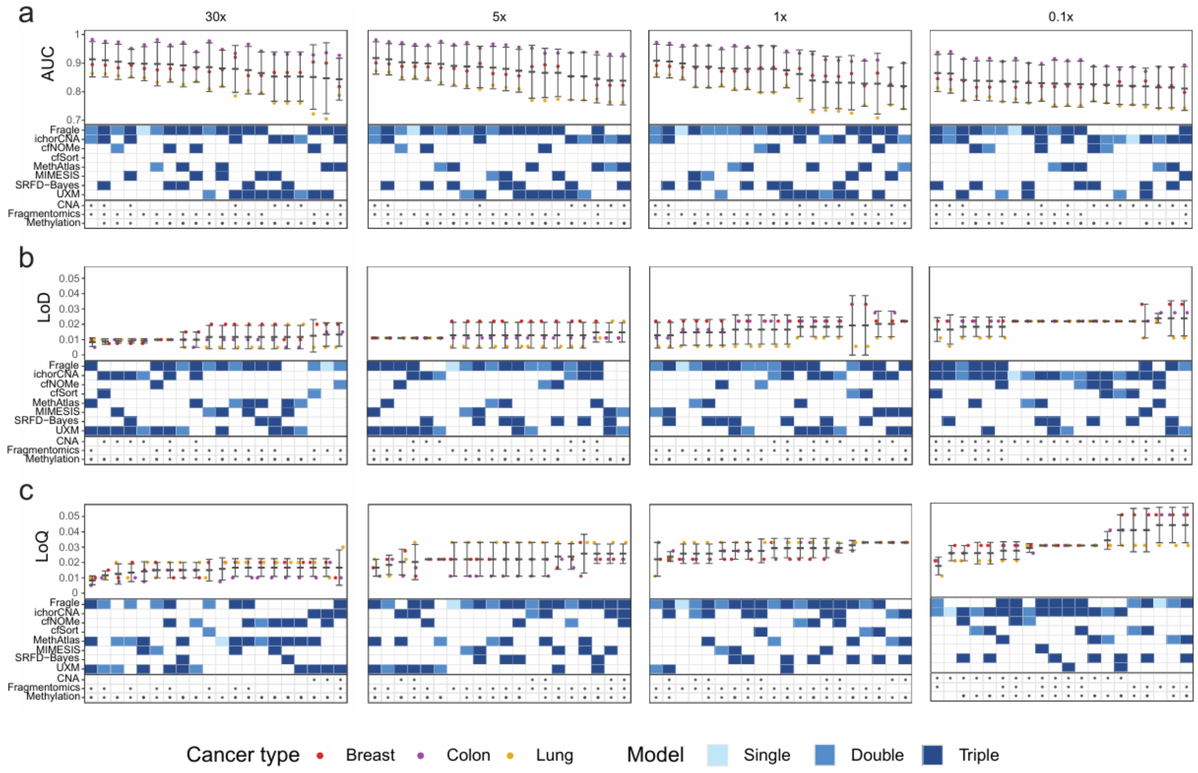


**Figure S8. Multi-method performance evaluation across sequencing depths.** a) Top-20 ranking of single-, double-, and triple-method ensembles (N = 92) for cancer versus healthy sample classification across distinct sequencing depths. Points represent the AUC for each cancer type, with bars indicating the mean and standard deviation. The feature modalities (CNA, Fragmentomics, Methylation) and method plurality (single, double, and triple) are indicated for each ensemble. b) Ranking of ensemble models based on their limit of detection (LoD) estimated from the sample dilution series. Points show the median LoD for each cancer type, with bars indicating the mean and standard deviation. c) Ranking of ensemble models based on their limit of quantification (LoQ) estimated from the sample dilution series. Points show the median LoQ for each cancer type, with bars indicating the mean and standard deviation.
